# Profiling Nucleotide Signalling Pathways and STING Agonist Activity with a Nucleotide Library

**DOI:** 10.1101/2025.11.11.687896

**Authors:** Indra Bekere, Rupert Öllinger, Marie Rose Schrimpf, Roland Rad, Carina C. de Oliveira Mann

**Affiliations:** Department of Bioscience, TUM School of Natural Sciences, Technical University of Munich, Garching, Germany; Institute of Molecular Oncology and Functional Genomics and Department of Medicine II, School of Medicine, Technical University of Munich, Munich, Germany; Gene Center and Department of Biochemistry, Ludwig-Maximilians-Universität München, Munich, Germany

## Abstract

Out of context dsDNA is detected by cGAS, which produces the cyclic dinucleotide 2′3′-cGAMP to activate STING and trigger downstream responses including transcriptional reprogramming, cell death and autophagy. STING agonists, among them non-hydrolysable cyclic dinucleotide analogues, are in active development to enhance immune activation in cancer treatment and vaccine adjuvants. Detailed knowledge of STING’s nucleotide preferences is critical for the development of effective therapeutics, yet the full spectrum of cyclic dinucleotides capable of activating STING has not been comprehensively defined. Especially considering the recent diversity of cyclic nucleotides identified across bacteria and invertebrates, the pool of potential STING agonists to be tested has expanded considerably. Here, we systematically dissect STING nucleotide preferences and distinguish STING-dependent from STING-independent nucleotide signaling pathways by treating THP-1 monocytes and pancreatic cancer cells with a nucleotide library. We identify a set of 27 cyclic dinucleotides that induce STING activation to different extents. Our data indicate that STING’s bias toward 2′3′-linked cyclic dinucleotides underlies its ability to also be activated by nucleotides containing purine–pyrimidine hybrids. In addition, we show that STING can be activated by diverse non-hydrolysable c-di-AMP and cGAMP isomers, thereby expanding the opportunities for designing STING agonists for therapeutic applications.

## Introduction

A very potent activator of innate immunity is out of context double-stranded DNA (dsDNA), which is sensed by the cGAS-STING pathway ^1^. dsDNA in the cytosol, e.g., from virus infection, damaged mitochondria or dying cells, binds to and activates a cyclic GMP-AMP synthase (cGAS) triggering synthesis of the cyclic dinucleotide (CDN) second messenger 2′-5′ / 3′-5′ cyclic GMP-AMP (2′3′-cGAMP). 2′3′-cGAMP in turn activates the receptor protein Stimulator of Interferon Genes (STING) at the endoplasmic reticulum (ER) membrane, which induces STING translocation to the Golgi and downstream signaling pathways, such as expression of type I interferons (IFN) and pro-inflammatory cytokines as well as cell death or autophagy.

Cyclic nucleotide second messengers are emerging as a chemically diverse class of bacterial signaling molecules, with recent studies expanding their chemical diversity to include varied phosphodiester linkages and both purine and pyrimidine compositions ^2–7^. Many of these CDNs are synthesized by cGAS-like enzymes and activate STING homologs across diverse organisms, where they function in antiviral defense analogous to the role of 2′3′-cGAMP signaling in humans. Notably, STING was first characterized as a receptor for the bacterial CDNs c-di-GMP, c-di-AMP and cGAMP before its endogenous metazoan ligand, 2′3′-cGAMP, was identified ^8–14^. This raises the possibility that additional, yet untested, bacterial CDNs may engage human STING and modulate its downstream signaling responses. Bacterial CDNs also operate in a wide range of pathways beyond antiviral defense, mediating diverse signaling functions outside the cGAS–STING axis, including roles in biofilm formation and metabolic regulation in bacteria, as well as differentiation in amoeba ^15,16^. Additional evidence for alternative CDN-mediated signaling pathways comes from studies in mice, where c-di-AMP not only activates STING but also signals through the cytosolic oxidoreductase RECON ^17^. This raises the question of whether humans harbor additional receptors or pathways dedicated to detecting cyclic oligo-nucleotides beyond STING-dependent 2′3′-cGAMP signaling.

From a therapeutic perspective, it is also important to determine the selectivity and potency of the signaling responses engaged by different cyclic nucleotides. The cGAS–cGAMP–STING axis is a key mediator of inflammation and several clinical applications targeting this pathway are being developed ^18^. cGAMP is rapidly emerging as a promising drug candidate as an endogenous small molecule, e.g., as a vaccine adjuvant or in cancer immunotherapy for induction of beneficial T-cell-driven inflammation. Therefore, STING agonists are being intensively developed as treatments in combination with immune checkpoint inhibitors to induce anti-tumor immunity ^19,20^. 2′3′-cGAMP is negatively charged rendering it non-cell permeable and dependent on specific transporters for uptake, which vary across cell types ^21–23^. 2′3′-cGAMP is also subjected to cleavage by extracellular enzymes ^24–26^. Ectonucleotide pyrophosphate phosphodiesterase 1 (ENPP1) is even exploited by cancer cells to degrade the immunotransmitter 2′3′-cGAMP and to generate immunosuppressive adenosine, thereby dampening anticancer immune responses within the tumor microenvironment ^27^. To overcome these obstacles, non-hydrolysable 2′3′-cGAMP analogues and synthetic STING agonists, including those inspired by bacterial CDN structures, are being developed as therapeutic strategies ^25,28^.

STING exhibits a binding and activation pattern that is highly selective for nucleotides with a specific nucleobase composition and phosphodiester linkage types ^29,30^. Although this topic has been the focus of extensive biochemical and structural investigations, there is still no comprehensive cellular approach, including the more recently discovered CDNs, to determine which molecular properties truly define STING activation and to what extent. Further adding to the complexity of STING agonist design, several STING single nucleotide polymorphisms (SNPs) are present in the human population, and these variants can markedly alter responsiveness to different CDNs ^10,31^. An in-depth understanding of STING’s nucleobase and linkage preferences, as well as how activation is influenced by CDN modifications that enhance stability, is essential for the rational design of effective STING agonists. Here, we comprehensively analyze STING-dependent and STING-independent nucleotide signaling pathways in human THP-1 cells. We identify a subset of CDNs that robustly activate STING and define the common structural features underlying STING’s nucleobase and linkage preferences. Notably, we also show that many bacterial CDNs fail to elicit any signaling in THP-1 monocytes, further underscoring the high specificity of CDN recognition and the strong selectivity of human STING for its endogenous ligand, 2′3′-cGAMP. In addition, we demonstrate how CDN modifications and their isomers, such as phosphorothioate-containing CDNs, that confer resistance to hydrolysis, modulate STING activation, providing insights with direct relevance for the development of therapeutic STING agonists.

## Results

### Nucleotide library screen reveals selective signaling by purine-containing cyclic dinucleotides

The nucleotide library was designed to encompass the broadest possible range of candidate signaling molecules, drawing on all known cyclic nucleotides and incorporating recently discovered bacterial cyclic oligonucleotides and their structural variants. We also included linear nucleotides, given that in humans linear 2′–5′ oligoadenylates function as antiviral second messengers by activating RNase L and inducing translational arrest ^32,33^. In addition, the linear guanosine tetraphosphate or pentaphosphate (p)ppGpp is a widespread bacterial second messenger involved in stress responses and has recently also been identified in metazoans ^34,35^. An additional rationale for their inclusion is that linear intermediates are generated during cyclic-nucleotide synthesis or degradation; yet it remains unknown whether these linear species possess signaling capacity in cells. The nucleotide library comprises 78 nucleotides, including 38 cyclic dinucleotides (CDNs), 9 cyclic oligonucleotides (CONs), 13 cyclic mononucleotides (CMNs) and 18 linear nucleotides (LNs) (Fig 1A, Table S1). CDNs include the natural STING ligand 2′3′-cGAMP, as well as a wide array of cyclic nucleotides that differ in nucleobase composition ranging from purine-containing and pyrimidine-containing species to purine–pyrimidine hybrids. They also encompass diverse structural diversity, including different combinations of 2′–5′ and 3′–5′ phosphodiester linkages, non-hydrolysable phosphorothioate analogues, and dideoxy cyclic nucleotides. These non-hydrolysable cyclic nucleotide compounds are resistant to hydrolysis by phosphodiesterases such as ENPP1 and are therefore more stable, inducing stronger and prolonged signaling. CONs include oligonucleotides ranging from three bases (e.g., cyclic tri-AMP) to six bases (e.g., cyclic hexa-AMP), several of which have been identified in bacteria and function in anti-phage defense systems ^7,36–38^. CONs also have the potential to signal in human cells as demonstrated by binding to and activation of mammalian reductase controlling NF-κB (RECON) by cyclic tri-AAG ^7^. CMNs include the well-known nucleotides in humans, 3′,5′-cGMP and 3′,5′-cAMP. cAMP was the first nucleotide second messenger to be discovered and regulates protein kinase A activation downstream of hormone and neurotransmitter signaling ^39,40^. Additional CMN diversity in the library is covered by nucleotides with 2′,3′ linkage isomers and alternative nucleobases, such as 3′,5′ xanthosine monophosphate (XMP), which has been identified in plants and mice ^41,42^. LNs in our library consist of adenosine or guanosine chains that range from two to five nucleobases in length. They include cleavage products of bacterial CDNs, such as linear 5′- phosphoguanylyl-(3′-5′)-guanosine (pGpG), a metabolite of c-di-GMP that may also function as an independent signaling molecule ^43,44^. Overall, our custom-made library comprises a diverse set of nucleotides, including molecules known to signal in human cells serving as controls as well as others whose existence or signaling capacity has not previously been established.

**Figure 1.**
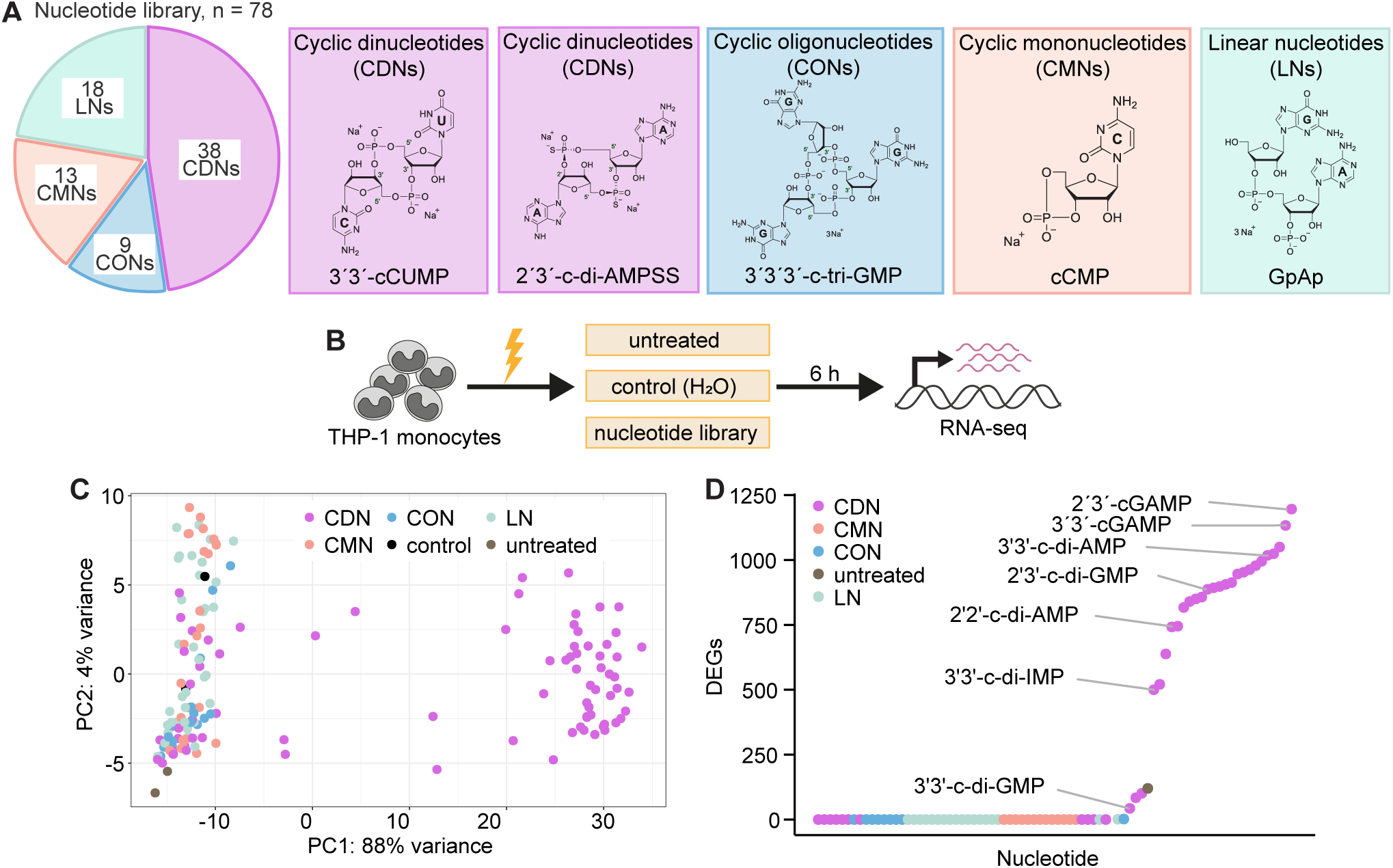
Analysis of nucleotide signaling pathways in THP-1 cells by RNA-seq (A) Composition of the nucleotide library with representative nucleotides shown. (B) Experimental setup. THP-1 monocytes were electroporated with 600 nM of each nucleotide in the library in duplicate or water as a control. Cells were harvested for RNA-seq analysis 6h after electroporation. Untreated cells without electroporation were used to monitor gene expression changes due to electroporation alone. (C) Principal component analysis of transcript counts from RNA-seq analysis showing all the analyzed samples and their respective nucleotide type. (D) Ranking of nucleotides by the number of differentially expressed genes (log2 fold change ≥ [1] and adjusted P-value < 0.05) when compared to control electroporation with water. CDN: cyclic dinucleotide, CMN: cyclic mononucleotide, CON: cyclic oligonucleotide, LN: linear nucleotide.

We monitored nucleotide-induced signaling in THP-1 monocytes, which express the STING HAQ (H72 A230 Q293) variant found in ca. 20% of human population. This variant can be activated by both the metazoan 2′3′-cGAMP and bacterial CDNs with 3′3′ linkage to induce a robust transcriptional response (Fig S1A) ^10,28,31^. THP-1 monocytes were electroporated with each nucleotide or with water as a control and harvested 6 h later for total transcriptome analysis by RNA-seq (Fig 1B). Untreated cells without electroporation were included as a control to assess the effects of electroporation alone, which induced only minimal changes in gene expression (Figs 1C, D). Electroporation ensured rapid delivery of nucleotides into cells minimizing degradation by extracellular phosphodiesterases ^24–26^. THP-1 monocytes were electroporated with 600 nM of each nucleotide, a concentration at which 2′3′-cGAMP induces robust STING activation and strong expression of *IFNB1* and *CXCL10* (Fig S1A). Principal component analysis (PCA) of RNA-seq transcript counts revealed that all CONs, LNs, CMNs and a subset of CDNs clustered with control and untreated cells, indicating that these nucleotides did not induce signaling (Fig 1C). The lack of signaling detected for the well-known CMNs cAMP and cGMP in humans may be explained by differences in treatment duration, which other gene-expression studies have shown to be critical for inducing a robust response. ^45,46^. In contrast, a distinct group of CDNs clustered separately from control samples, consistent with induction of gene expression changes. Analysis of differentially expressed genes (DEGs), comparing each nucleotide to the control, showed that the endogenous STING ligand 2′3′-cGAMP induced the strongest transcriptional response, with a total of 1,196 DEGs (Fig 1D, Table S1). The ten nucleotides inducing the greatest number of DEGs were in fact hydrolysable and non-hydrolysable isomers of cGAMP and c-di-AMP, both of which are established STING agonists. Overall, gene expression changes were induced by 27 CDNs in our library, all of which were composed of purine nucleobases guanosine, adenosine or inosine and their hydrolysable and non-hydrolysable isomers (Fig S1B). Only 3 DEGs in total were identified for non-CDN nucleotides cyclic tetra-GMP and ApA, which belong to the CONs and LNs class, respectively. Taken together, the results of our nucleotide library screen demonstrate that nucleotide signaling is highly specific, with many compounds failing to elicit any response, while simultaneously revealing that STING displays strong selectivity toward a narrow subset of purine-containing CDNs that activate signaling in THP-1 cells.

### A common STING-mediated transcriptional signature is induced by distinct CDNs

We next asked whether the responses induced by different CDN isomers were identical, or whether variations in nucleobase composition or phosphodiester linkage resulted in distinct transcriptional outcomes. Several structures of STING bound to different CDNs, including bacterial cGAMP, c-di-AMP and c-di-GMP, have shown that CDNs can induce distinct STING conformations, suggesting that ligand-specific structural states may influence the magnitude or even type of downstream signaling responses ^29,30,47,48^. In addition, we sought to determine whether individual CDNs elicit responses beyond canonical STING signaling, as exemplified by c-di-AMP in mice, which can also engage the receptor RECON ^17^. Thus, we further analyzed the gene expression signatures induced by the signaling CDNs and compared them to those elicited by 2′3′-cGAMP, which induces exclusively STING-dependent transcriptional changes in THP-1 cells ^49^. Clustering of 2561 unique DEGs, obtained from pooling all DEGs induced by 27 signaling CDNs, revealed two major clusters (Fig 2A). The “upregulated” cluster comprised genes whose expression increased in response to CDN treatment relative to control cells, whereas the “downregulated” cluster contained genes whose expression decreased. All signaling CDNs induced a similar transcriptional signature across both clusters, with variation primarily in magnitude: 3′3′-c-di-GMP induced the weakest changes, while 2′3′-cGAMP triggered the largest number of DEGs and some of the strongest transcriptional responses.

**Figure 2.**
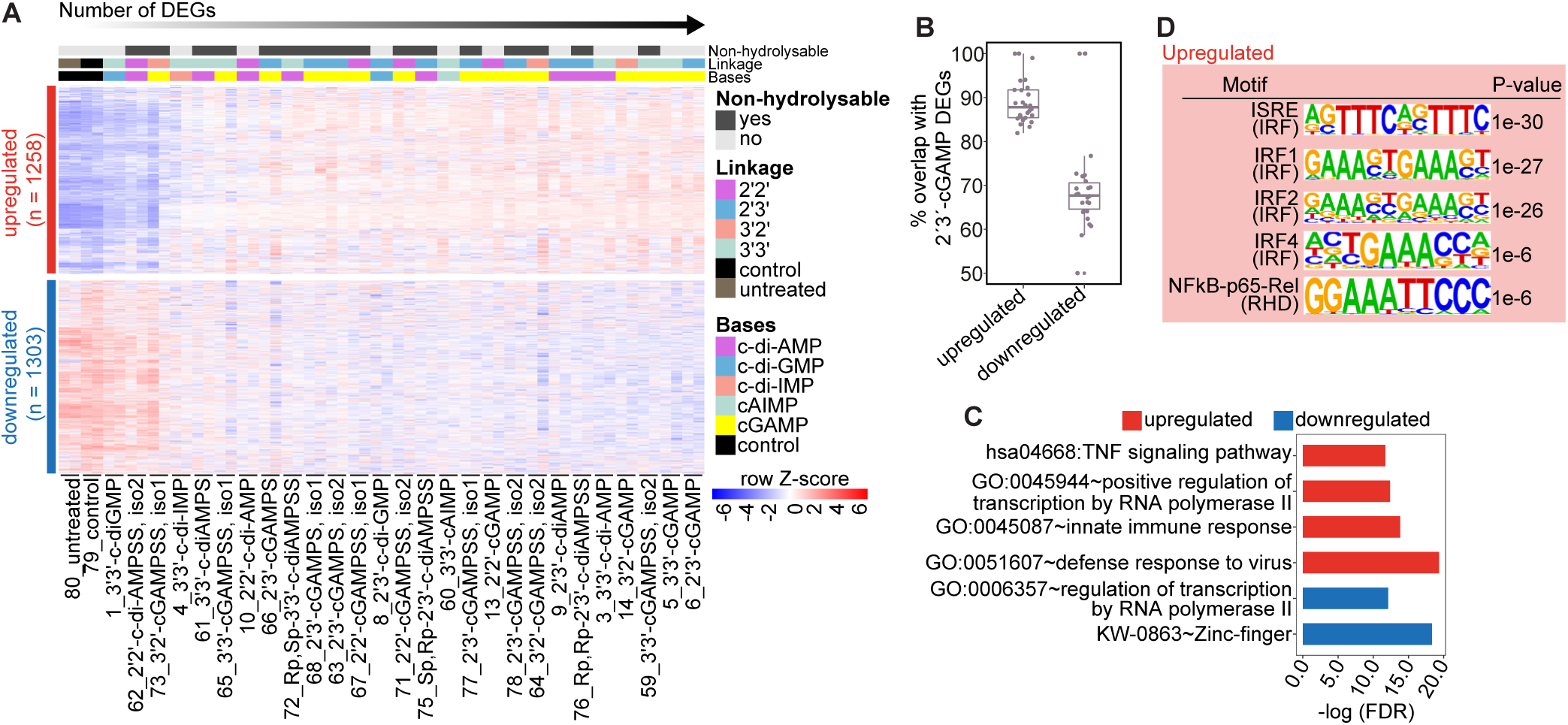
Characterization of gene expression signature due to signaling by CDNs (A) Heatmap showing clustering of all DEGs pooled from signaling 27 CDNs, which identified two clusters. Treatments are sorted by the number of DEGs that the nucleotides induced. Normalized transcript counts from variance stabilizing transformation (vst) were used, counts are row scaled. (B) Boxplot showing percentage of upregulated and downregulated DEGs for each signaling CDN (dots) that overlap and were identified as DEGs also for 2′3′-cGAMP. Boxes encompass the twenty-fifth to seventy-fifth percentile changes. The central horizontal bar indicates the median. (C) Pathway enrichment analysis for the “upregulated” and “downregulated” genes from (B). (D) Transcription factor motif enrichment analysis for the genes from “upregulated” cluster in (B).

This indicates that the CDNs induce responses of varying strengths that closely resemble those triggered by 2′3′-cGAMP, consistent with STING-dependent signaling. Indeed, when analyzing each CDN individually, approximately 90% of upregulated DEGs and 65% of downregulated DEGs were shared with those induced by the endogenous STING agonist 2′3′ -cGAMP (Fig 2B).

To further exclude any STING-independent signaling or transcriptional responses, we analyzed the DEGs induced by all signaling CDNs but not by 2′3′-cGAMP by comparing the overlaps of upregulated and downregulated genes. Pooling the responses of all signaling CDNs uncovered 536 upregulated and 829 downregulated DEGs that were not significantly regulated by 2′3′-cGAMP. (Fig S2A). However, clustering analysis showed that these genes still followed a pattern of regulation similar to that induced by 2′3′-cGAMP, despite not passing the cut-off of log_2_ fold change of [1] and adjusted *P-value* of 0.05 thresholds used to define DEGs (Fig S2B). Thus, signaling CDNs converge on a common STING-dependent transcriptional response. The induced genes were strongly enriched for innate immune pathways related to anti-viral defense, including interferon-stimulated genes (ISGs) and NF-κB–driven TNF signaling (Fig 2C). Induction of interferon signaling and the NF-κB pathways is a hallmark of STING activation, which is also reflected in the strong enrichment of ISRE and IRF transcription factor motifs, as well as NF-κB motifs, among the upregulated DEGs (Fig 2D). Furthermore, both upregulated and downregulated genes were enriched for pathways associated with transcriptional regulation, comprising numerous transcription factors from diverse families (Fig 2C, Fig S2C). The upregulated genes included basic leucine zipper domain superfamily members such as JUNB, FOSL1, FOS, ATF3, CEBPB and STAT proteins, all of which are well-established regulators of inflammatory responses ^50,51^. In contrast, the downregulated genes were significantly enriched for Zinc finger (ZNF) transcription factors containing a Krüppel associated box (KRAB) repressor domain (Fig S2C) ^52^. Collectively, our analysis identified a set of 27 CDNs that induce a spectrum of STING activation, modulating inflammatory and transcriptional pathways.

### STING activation is skewed towards cyclic dinucleotides with 2′3′ phosphodiester linkage

Since no additional signaling was observed from nucleotide compounds other than the CDNs that activate STING and trigger the canonical 2′3′-cGAMP response, albeit with different magnitudes, we next sought to use these findings to identify the shared features that determine STING’s nucleotide activation preferences in THP-1 cells, which carry the STING HAQ variant^48^. Multiple STING single nucleotide polymorphisms (SNPs) exist in the human population, influencing responsiveness to different CDNs. Both human STING WT and HAQ variants respond to CDNs with a 2′3′ linkage as well as bacterial CDNs with a 3′3′ linkage, whereas the R232H variant is unresponsive to CDNs containing a 3′3′ linkage ^10,28,31^. Our analysis in THP-1 cells demonstrated STING-dependent signaling for 27 out of 38 CDNs included in our nucleotide library. Notably, among CDNs with 3′3′ phosphodiester linkages, STING activation was observed exclusively for molecules composed of purine, but not pyrimidine, nucleobases. (Fig 3A). We further examined STING’s preference for specific phosphodiester linkage types using purine-containing cGAMP, c-di-AMP and c-di-GMP isomers. All 2′2′, 2′3′, 3′3′ or 3′2′ (cGAMP only) linkage isomers of c-di-AMP and cGAMP induced high levels of STING activation (Fig 3B). Across the analyzed isomers, STING displayed a strong preference for 2′3′ linkages. Both 2′3′-c-di-GMP and 2′3′-c-di-AMP induced robust activation comparable to 2′3′-cGAMP (Fig 3B), whereas 3′3′ c-di-GMP acted only as a weak agonist and 2′2′ c-di-GMP showed no activity. Notably, we identified 2′2′ c-di-AMP as a previously unrecognized STING agonist, thereby expanding the repertoire of known activating CDNs. The weak activity of 3′3′-c-di-GMP is consistent with earlier reports in mouse and human STING ^14,28,30,47,48^ and is thought to result from its higher affinity, which stabilizes a more open STING conformation and promotes cooperative activation rather than the closed-state polymerization induced by 2′3′-linked ligands and 3′3′-c-di-AMP ^30,47,48^.

**Figure 3.**
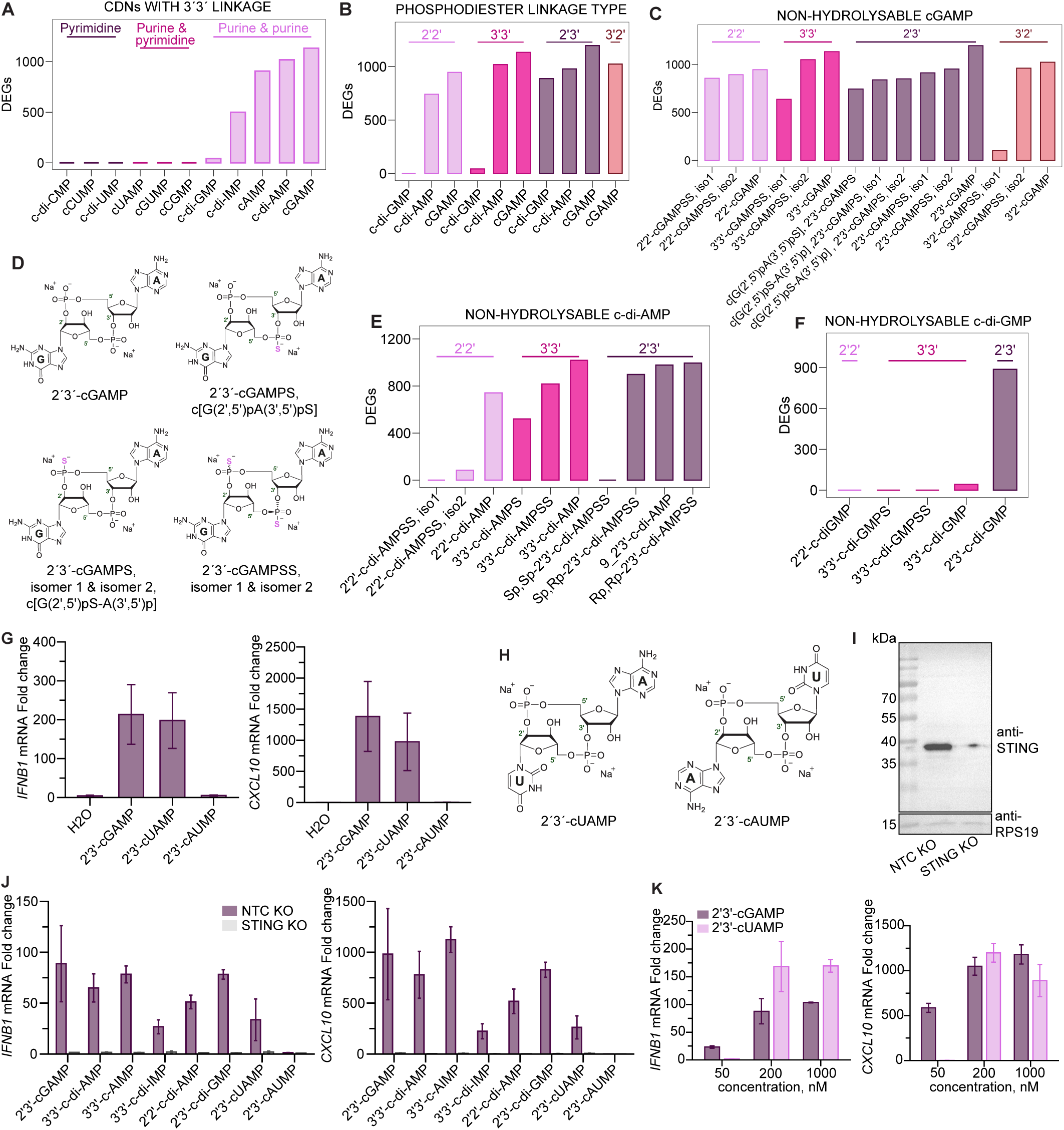
Analysis of preferred CDN features for STING activation (A) Bar plot showing number of DEGs for all nucleotides in the library with 3′3′ phosphodiester linkage grouped by purine and/ or pyrimidine base composition. (B) Bar plot showing number of DEGs for all c-di-GMP, c-di-AMP and cGAMP isomers in the library grouped by the phosphodiester linkage type. (C) Bar plot showing number of DEGs for all hydrolysable and non-hydrolysable cGAMP isomers in the library. (D) Depiction of different cGAMP isomers containing thiophosphate modifications at different positions. (E) and (F) Bar plots showing number of DEGs for all hydrolysable and non-hydrolysable c-di-AMP (E) and c-di-GMP (F) isomers in the library. (G) RT-qPCR analysis of *IFNB1* and *CXCL10* expression after electroporation of THP-1 monocytes with 600 nM 2′3′-cGAMP, 2′3′-cUAMP, 2′3′-cAUMP or water (control) for 6 h. (H) Depiction of 2′3′-cUAMP and 2′3′-cAUMP. (I) Western Blot analysis showing STING levels in THP-1 non-targeting control (NTC) or STING KO cells. (J) RT-qPCR analysis of *IFNB1* and *CXCL10* expression after electroporation of THP-1 NTC KO or STING KO cells with 600 nM of indicated nucleotides or water as a control for 6 h. (K) RT-qPCR analysis of *IFNB1* and *CXCL10* expression after electroporation of THP-1 monocytes with 50 nM, 200 nM or 1000 nM of 2′3′-cGAMP or 2′3′-cUAMP for 6 h. Bars represent means from at least two replicates and error bars represent standard deviation.

Our nucleotide library includes several phosphorothioate-containing isomers of c-GAMP, c-di-GMP and c-di-AMP, in which a non-bridging oxygen on one or both phosphates is substituted with a sulfur atom (Fig 3D). This modification confers resistance to phosphodiesterase-mediated cleavage, making these analogues more attractive candidates for immunomodulatory applications ^25,28^. Among the cGAMP isomers, all non-hydrolysable linkage isomers produced robust STING activation, with the exception of the 3′2′-cGAMPSS isomer 1 (Fig 3C). For the 2′3′-cGAMP analogues, signaling strength was similar whether sulfur was incorporated at both phosphates or restricted to the 2′–5′ or 3′–5′ linkage (Figs 3C, D), although these phosphorothioate forms generally induced slightly lower activation than unmodified 2′3′-cGAMP. In contrast, the behavior of other phosphorothioate-modified cyclic dinucleotides varied by phosphodiester linkage type. While natural 2′2′-c-di-AMP activated STING, introducing sulfur atoms at both phosphates eliminated its activity (Fig 3E). Non-hydrolysable 3′3′-c-di-AMP analogues retained STING activation capacity, albeit at reduced levels compared with unmodified 3′3′-c-di-AMP. Conversely, thiophosphate modification of 3′3′-c-di-GMP completely abolished signaling (Figs 3E, F). The ability of 2′3′-c-di-AMP to induce STING signaling was highly dependent on the stereochemistry of the thiophosphate groups. The Rp,Rp and Sp,Rp dithio-substituted diastereomers triggered strong STING activation, whereas the Sp,Sp form did not elicit signaling (Fig 3E). In fact, the Rp, Rp-2′3′c-di-AMP analogue, also known as ADU-S100 and MIW815, has progressed into clinical trials for advanced and metastatic solid tumors as well as lymphomas ^19,20^.

In summary, our analysis of STING responses to 38 distinct CDNs revealed a strong preference for 2′3′ phosphodiester linkages, which outweighed the influence of base composition. Even weaker agonists, such as 3′3′ c-di-GMP, exhibited markedly enhanced signaling when configured with a 2′3′ linkage. In addition, we identified several non-hydrolysable c-di-AMP and cGAMP isomers that induced robust STING activation, highlighting candidates with improved stability and prolonged signaling potential for therapeutic applications.

Recent studies have expanded the diversity of known CDNs as well as cGAS-like enzymes and STING homologs in bacteria, Drosophila and metazoans ^2–5,7^. Of note, STING homologs in coral *S. pistillata* can be activated not only by 2′3′-cGAMP but also by purine-pyrimidine CDN 2′3′-cUAMP, which contains a 2′3′ phosphodiester linkage ^4^. Structural studies further demonstrated that 2′3′-cUAMP binds both to *S. pistillata* and human STING and induces a closed conformation similar to that triggered by 2′3′-cGAMP ^4,30,47^. The strong preference of human STING for 2′3′-linked CDNs (Fig. 3B), together with the ability of 2′3′-cUAMP to induce a closed, active STING conformation (Li et al., 2023), suggests that human STING can also be activated by pyrimidine-containing 2′3′ linked CDNs ^4^. Indeed, electroporation of THP-1 monocytes with 2′3′-cUAMP resulted in robust induction of *IFNB1* and *CXCL10* expression, with levels comparable to those elicited by 2′3′-cGAMP (Figs 3G, H). In contrast, 2′3′-cAUMP failed to induce signaling, indicating a requirement for both the specific pyrimidine base position and the 2′3′ phosphodiester linkage. To confirm the dependence on STING, we generated STING-KO THP-1 cells and verified that the induction of *IFNB1* and *CXCL10* by 2′3′-cUAMP, as well as by the other signaling CDNs in our screen (Fig 2), was abolished in the absence of STING (Fig 3I, J). Dose-response analysis of 2′3′-UAMP and 2′3′-cGAMP further showed that 2′3′-cGAMP activated STING more efficiently at lower concentrations, although both CDNs achieved similar activation at higher concentrations (Fig 3K). Taken together, our data reveal that STING can tolerate pyrimidine bases at a specific position for activation and signaling when presented within its preferred 2′3′ phosphodiester linkage.

### Potent STING agonists drive cell death in pancreatic cancer cells

STING agonists are currently being developed for cancer therapy to induce protective antitumor immunity as well as cancer cell death ^19,20,28,53,54^. The consequences of STING activation in tumors are multifaceted and shaped by both intratumoral heterogeneity and the surrounding microenvironment ^55,56^. On the one hand, STING signaling can exert adverse effects by promoting T cell death, thereby generating an immunosuppressive, tumor-supportive microenvironment ^57–60^. On the other hand, STING activation can be advantageous by eliciting inflammatory responses within the tumor microenvironment and inducing interferon-dependent cell-cycle arrest in cancer cells, as shown in pancreatic ductal adenocarcinoma (PDAC) models ^61,62^. Thus, because STING activation does not necessarily correlate with antitumor activity, we leveraged our extensive nucleotide library, together with the distinct response patterns observed in THP-1 cells, to define the spectrum of nucleotides capable of activating STING or other receptors and inducing cell death in pancreatic cancer cells. To this end, we electroporated the nucleotide library into the human pancreatic cancer cell line DANG, which responds efficiently to interferon stimulation leading to cell cycle arrest ^61^. Cell death was monitored by time-lapse live-cell imaging using the cell-death dye YOYO-3 over 70 hours (Fig 4A). Consistent with our findings in THP-1 cells, induction of cell death was observed exclusively for purine-containing CDNs (Fig 4B, Fig S3). The most potent inducers of cell death were the non-hydrolysable isomers of cGAMP and c-di-AMP, as well as 2′3′-c-di-AMP, 2′3′-c-di-GMP and endogenous 2′3′-cGAMP, demonstrating STING’s preference towards the 2′3′ also in pancreatic cancer cells (Fig 4B, Fig S3). The non-hydrolysable CDN isomers induced the highest levels of cell death when measured 60 hours after electroporation, consistent with a cellular effect driven by their increased stability and resistance to phosphodiesterase-mediated cleavage (Fig 4C, D). Electroporation of 2′3′-cGAMP also induced *Ifnb1* expression in mouse pancreatic cancer cells (Fig 4E), indicating the potential for beneficial STING signaling across pancreatic cancer models in different organisms. Collectively, our nucleotide-library experiments highlight the remarkable specificity of nucleotide signaling in diverse cell types and demonstrate how the signaling capacity of distinct CDNs can be translated across these cellular contexts and responses.

**Figure 4.**
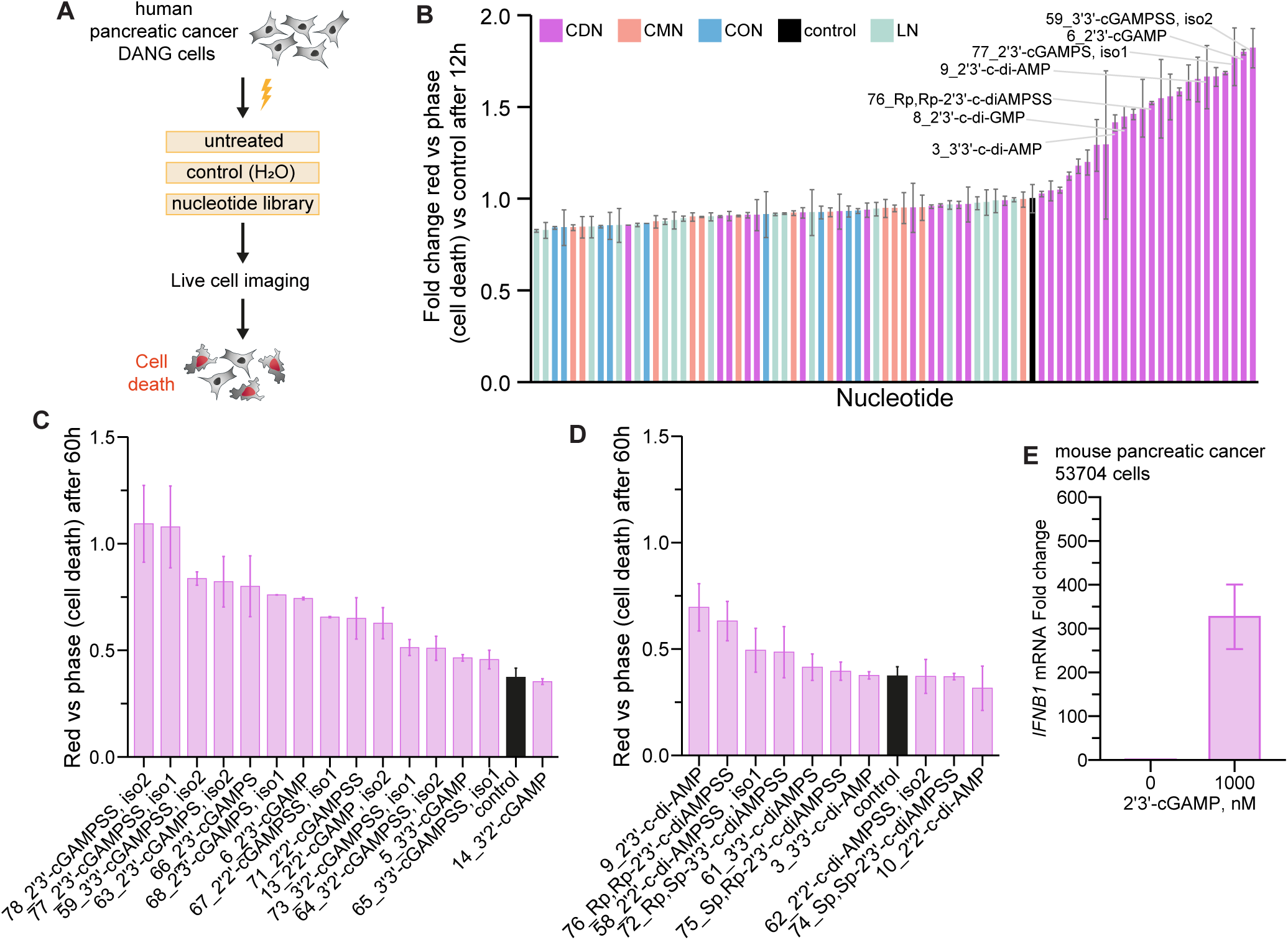
Analysis of cell death in pancreatic cancer cells (A) Experimental setup. DANG pancreatic cancer cells were electroporated with 2 μM of each nucleotide form the library in duplicate or water as a control. Cell death was monitored by live cell imaging over 70 h in the presence of red fluorescent cell death dye YOYO-3. (B) Ranking of nucleotides by the level of induced cell death (red vs phase signal) in DANG cells when compared to control electroporation with water at 12h post-treatment. (C) and (D) Ranking of cGAMP (C) and c-di-AMP (D) isomers from the library for induction of cell death in DANG cells at 60 h post-treatment. (E) RT-qPCR analysis of *IFNB1* expression after electroporation of mouse pancreatic cancer cells with 1000 nM of 2′3′-cGAMP for 6 h. CDN: cyclic dinucleotide, CMN: cyclic mononucleotide, CON: cyclic oligonucleotide, LN: linear nucleotide. Bars represent means from at least two replicates and error bars represent standard deviation.

## Discussion

As the diversity of nucleotide second messengers continues to expand, particularly in bacteria, it has become increasingly important to determine whether these molecules can elicit signaling in human cells either through alternative pathways or via STING. Given that human STING responds to both endogenous metazoan 2′3′-cGAMP and selected bacterial CDNs, we systematically evaluated whether other nucleotide second messengers can activate STING in human cells. Previous nucleotide screens investigating STING signaling have used diverse human and mouse cell types expressing different STING variants and have monitored distinct cellular responses, making it difficult to analyze and compare STING activation across studies. ^28,63–66^. Moreover, purine-containing CDNs have traditionally been the primary focus when analyzing STING agonists. Using our comprehensive nucleotide library, we evaluated the ability of diverse CDNs from various organisms, including those containing pyrimidine bases as well as other cyclic nucleotides such as cyclic mononucleotides and oligonucleotides, to activate endogenous STING in THP-1 cells and pancreatic cancer cells. We observed a striking specificity of nucleotide responses, identifying distinct sets of nucleotides that induced signaling (27 CDNs in total) and others that failed to do so, underscoring the remarkable selectivity that cyclic nucleotide receptors exhibit toward their respective ligands. Overall, our findings support the idea that additional nucleotide receptors beyond STING, with similarly high specificity, may be uncovered by monitoring nucleotide-mediated signaling in other cell types and experimental contexts.

In line with our data, the purine-containing CDNs were reported to be the most potent STING agonists in other nucleotide screens with different cellular models and readouts: i) in suppression of virus infection with SARS-CoV-2, CHIKV, WNV, and ZIKV in human fibroblasts (HFF-1) and Calu-3 cells ^63,64^; ii) in induction of IRF3 reporter activation in the mouse macrophage cell line analyzing 3′3′ CDNs composed of A, G, C, and U. ^65^ and iii) in interferon reporter activation in HEK293T cells expressing the major STING variants and analyzing inosine-containing CDNs in combination with with A, G, C, and U nucleobases, across 2′3′, 2′2′, and 3′3′ linkages ^66^. Analysis of pyrimidine-containing nucleotides has focused primarily on CDNs with 3′3′ linkages, which we show are not STING’s preferred linkage type. Indeed, in the context of 2′3′ and 2′2′ linkages, robust STING activation was observed for the mixed purine–pyrimidine CDNs 2′2′-cUIMP and 2′3′-cUIMP ^66^. This study, together with our observation that human STING is activated by the pyrimidine-containing CDN 2′3′-cUAMP, challenges the previously proposed strict preference for purine-containing CDNs. It indicates that combinations of different linkage types broaden the repertoire of CDNs capable of activating STING to include pyrimidine-containing nucleotides. Signaling by pyrimidine-containing CDNs introduces an exciting new dimension to the STING research field, particularly regarding the development of non-hydrolysable analogues for therapeutic use and the characterization of their cellular transport and degradation. ^21–26^.

Many of the CDNs and STING activators in our library belong to a rapidly expanding list of bacterial CDNs that play important roles in bacterial homeostasis, virulence and anti-phage defense ^67,68^. STING activation by bacterial CDNs is critical for mounting protective innate immune responses during infection of Gram-positive bacteria *Listeria* and Gram-negative *Chlamydia trachomatis* ^13,69,70^. These findings suggest that sensing bacterial CDNs or their degradation products represents a valuable strategy for detecting invading bacteria, either through STING or additional nucleotide-sensing receptors ^71,72^. For instance, STING activation may be particularly relevant in the gut, for sensing multiple microbiota-derived CDNs, in addition to detecting pathogenic disruption of the epithelial barrier ^73–75^. Our investigation is limited to the THP-1 monocyte and PDAC cell lines, and we cannot exclude the possibility that these nucleotides may signal in other human cell types, particularly if alternative receptors, other than STING, are expressed differentially across tissues and not present in THP-1s.

Our analysis reveals a broad set of STING agonists that induce activation to varying degrees, which likely corresponds to differences in ligand binding strength and their ability to drive STING dimer closure. High-affinity ligands such as 2′3′-cGAMP and 3′3′ c-di-AMP engage deeply within the binding pocket and induce a closed dimer conformation. The 2′3′ linkage imposes unique structural constraints that promote this closed conformation leading to STING polymerization and downstream activation ^4,30,47,48^. In contrast, the earliest described bacterial STING agonist, 3′3′ c-di-GMP, functions as a weaker activator both in cells and in vitro, driving a more open, apo-like STING conformation and an alternative cooperative mode of activation^28,30,47,63,64^.

Downstream signaling, in particular, is a critical aspect to consider in the development of effective STING agonists. Despite promising preclinical studies demonstrating the induction of antitumor immunity, most developed STING agonists have failed to translate these effects in clinical trials ^53,76^. For instance, the non-hydrolysable 2′3′-c-di-AMSS (Rp,Rp) is under clinical development for the treatment of metastatic, solid tumors or lymphomas, but has so far shown limited efficacy in clinical trials ^19,20^. Additional strategies are being explored ^77^ and may include gene therapies to deliver enzymes in cells for synthesis of STING-activating ligands ^76^. Other approaches involve modifications of STING agonists including altering the position of thiophosphate substitution ^78,79^, modifying phosphodiester linkages ^80^, incorporating locked nucleic acids ^81^ or dideoxy derivatives ^82^, designing sugar-modified analogues ^83^, inosine-containing CDNs ^66,84^ or non-nucleotide based agonists ^85–87^. Collectively, these studies together with our findings demonstrate that the STING ligand-binding pocket can accommodate a broad range of hydrolysable, non-hydrolysable, and chemically modified CDNs, providing multiple avenues for the development of more effective STING agonists. And regarding the discovery of novel nucleotide-based second messnegr signaling pathways in humans, our approach reveals that we could be missing the receptor or that our nucleotide version is not corresponding the human version and thus understanding uncharacterized nucleotidyltransferases and enzymes synthezising these signals might be the more promising avenue. With respect to the discovery of novel nucleotide-based second-messenger pathways in humans, our inability to identify additional signaling activities likely reflects either the absence of the relevant receptor in the cell lines we screened or the possibility that the nucleotide variants we tested do not match the true endogenous human signals. Thus, investigating the activation mechanisms of uncharacterized nucleotidyltransferases and other enzymes that generate these nucleotide second messengers offers an additional path forward.

## Materials and Methods

### Cell lines

THP-1 wild type, THP-1 NTC KO (this study) and THP-1 STING KO (this study) cells were cultured in suspension in RPMI supplemented with 10% FBS at 37°C in 5% CO_2_ and passaged at dilution 1:10. THP-1 cells were a kind gift from Prof. Andreas Pichlmair (TUM, Munich). Pancreatic cancer cell lines DANG and mouse pancreatic cancer (mPC) 53704 were a kind gift from Prof. Roland Rad (TUM, Munich). HEK293T (ATCC Cat#CRL-11268), DANG and mPC 53074 were cultured in DMEM supplemented with 10% FBS at 37°C in 5% CO_2_ and passaged at a dilution 1:10 by washing with PBS and detached with 0.25% trypsin. All cell lines were regularly tested negative for mycoplasma contamination using TaKaRA PCR Mycoplasma Detection Set (TaKaRa).

### Generation of cell lines

Two different gRNA sequences for STING and non-targeting control (NTC) knockout were cloned in pLentiCRISPRv2 vector (Addgene #52961 ^88^) with puromycin resistance. Guide RNA sequences were STING gRNA1 CGGTCGGCCCGCCCTTCACT, STING gRNA2 GGGAATTTCAACGTGGCCCA, NTC gRNA1 CGACGGAGGCTAAGCGTCGCAA, NTC gRNA2 CGCGCTTCCGCGGCCCGTTCAA. Primers encoding target guide RNA sequences were annealed and used for ligation with linearized vector from restriction digest with BsmbI enzyme. Successful cloning was verified by sequencing and target vectors were further used for lentivirus generation.

Lentiviral transductions were used to generate THP-1 STING KO and NTC KO cell lines. For knockout generation lentiviral particles were generated by transfecting HEK293T cells with psPAX2 ^89^ and pMD2.G-VSV-G ^89^ packaging plasmids together with pLentiCRISPRv2 vector with puromycin resistance encoding *S. pyogenes* CRISPR-Cas9 and two different gRNA sequences. psPAX2 and pMD2.G-VSV-G plasmids were kindly provided by Prof. Andreas Pichlmair (TUM, Munich). Lentiviral particles were harvested 48h post transfection and used to transduce THP-1 cells followed by selection with 1.5 µg/ml puromycin one day post infection. Selection was terminated once there were no more viable control untransduced cells with puromycin selection. Successful KO was validated by WB analysis.

### Western Blot analysis

For validation of STING KO in THP-1 cells, cells were lysed in NP-40 lysis buffer (50mM Tris-HCl pH 7.5, 150mM NaCl, 1% NP-40, 5mM EDTA) supplemented with 1X Complete protease inhibitor (Sigma-Aldrich) for 20-30min on ice. Soluble fraction was separated by centrifugation for 10min, 21’000xg, 4°C, mixed with 1x Laemmli sample buffer and boiled for 5-10min at 95°C. Protein was resolved by 12% SDS-PAGE and transferred to 0.45 µm PVDF membrane. Membranes were blocked in 5% non-fat dry milk, 0.1% Tween-20 in PBS and incubated with the following primary antibodies: anti-STING (Cell Signaling, #13647, 1:1000 dilution), anti-RPS19 (ThermoFisher, #A304-002A, 1:2000 dilution). Afterwards membranes were probed with HRP-conjugated secondary antibodies goat anti-rabbit IgG (Dako, P0448). Immunoblots with HRP signal were developed with the SuperSignal West Femto kit (Thermo Fisher Scientific) and imaged with the Bio-Rad ChemiDoc Imaging System.

### Electroporation of nucleotides

The nucleotide library consists of compounds synthesized and quality-controlled by Biolog LSI. All nucleotides are commercially available from Biolog LSI, and their corresponding catalogue numbers are provided in Table S1.

Electroporation was performed with Neon Transfection kit (Thermo Fisher Scientific) following manufacturer’s instructions. THP-1 cells were collected and washed with PBS and resuspended in Buffer R to a final density of 5×10^6^ cells /ml with 600 nM of nucleotides if not indicated otherwise. For electroporation 5×10^5^ cells were used with 100 μl Neon tip and electroporated with pulse voltage 1400 V, pulse width 20 ms and pulse number 2 in 3 ml E2 Electrolyte Buffer. Afterwards cells were transferred to 1ml pre-warmed media in a 24 well plate and incubated at 37°C in 5% CO_2_ for 6 h until harvest.

DANG cells were detached, washed with PBS and resuspended in Buffer R to a final density of 1.5×10^6^ cells /ml with 2 μM of nucleotides. Cells were electroporated with 10 μl tip with pulse voltage 1400 V, pulse width 20 ms and pulse number 1 in 3 ml E Electrolyte Buffer. Afterwards cells were transferred to a well in a 96-well plate filled with DMEM, 10 % FBS and 250 nM YOYO-3 cell death dye. Cell death was monitored using live cell imaging system Incucyte S3 (Sartorius) over 70 h with scans every 2 h. Cell death was quantified as red signal vs phase. For electroporation of mPC 53074 cells, cells were washed with PBS and resuspended in Buffer R to a final density of 6×10^6^ cells /ml with 1 μM of nucleotides. Cells were electroporated with 100 μl tip with pulse voltage 1400 V, pulse width 20 ms and pulse number 2 in 3 ml E2 Electrolyte Buffer. Afterwards cells were transferred to a well in a 12-well plate filled with 1ml DMEM, 10 % FBS for 6 h until harvest.

### RT-qPCR analysis

Total RNA was extracted using the NucleoSpin RNA Plus kit (Macherey-Nagel) according to the manufacturers’ protocol. Total RNA was used for reverse transcription with PrimeScript RT reagent Kit with gDNA Eraser (TaKaRa) according to the manufacturers’ instructions. Relative transcript quantification was obtained by qPCR with the transcript-specific primers using PowerUp SYBR Green master mix (Thermo Fisher) on a QuantStudio3 PCR system (Thermo Fisher). Ct values were obtained using the QuantStudio software and averaged across technical replicates. The transcript levels were normalized to the levels of a housekeeping gene GAPDH. The oligonucleotides used for the analysis in THP-1 cells were: *IFNB1* forward ACGCCGCATTGACCATCTAT, *IFNB1* reverse GTCTCATTCCAGCCAGTGCTA, *CXCl10* forward AAGTGGCATTCAAGGAGTACCT, *CXCL10*reverse GGACAAAATTGGCTTGCAGGA, *GAPDH* forward GATTCCACCCATGGCAAATTC, *GAPDH*reverse AGCATCGCCCCACTTGATT. The oligonucleotides used for the analysis in mouse pancreatic cancer 53074 cells were: *RPLP0* forward GGATCTGCTGCATCTGCTTG, *RPLP0* reverse GCGACCTGGAAGTCCAACTA, *IFNB* forward CGGAGAAGATGCAGAAGAGT, *IFNB* reverse TCAAGTGGAGAGCAGTTGAG.

### RNA-seq library preparation, sequencing and data processing

Library preparation for bulk-sequencing of poly(A)-RNA was done as described previously ^90^. Barcoded cDNA of each sample was generated with a Maxima RT polymerase (Thermo Fisher) using oligo-dT primer containing barcodes, unique molecular identifiers (UMIs) and an adaptor. 5′-Ends of the cDNAs were extended by a template switch oligo (TSO) and full-length cDNA was amplified with primers binding to the TSO-site and the adaptor. NEB UltraII FS kit was used to fragment cDNA. After end repair and A-tailing a TruSeq adapter was ligated and 3′-end-fragments were finally amplified using primers with Illumina P5 and P7 overhangs. The library was sequenced on a NextSeq1000 (Illumina) with 65 cycles for the cDNA in read1 and 19 cycles for the barcodes and UMIs in read2. Data was processed using the published Drop-seq pipeline (v1.12) to generate sample- and gene-wise UMI tables ^91^. Reference genome (GRCh38) was used for alignment. Transcript and gene definitions were used according to GENCODE v38.

Raw counts were used for differential expression analysis with DESeq2 (v1.42.1) package ^92^. For plotting of PCA plots and heatmaps counts were transformed using variance-stabilized transformation. Differentially expressed genes were defined as log2 fold change ≤ −1 or ≥ +1 and adjusted P value < 0.05. Pathway analysis was performed using DAVID analysis tool ^93^.

### TF motif analysis

TF motif enrichment for known motifs was performed using HOMER package ^94^. Command *findMotifs.pl* was used and a list of gene symbols was supplied as an input. Motifs were searched in the region 400 bp upstream and 100 bp downstream of the TSS by specifying parameters *-start −400 -end 100*. For presentation of enriched TF motifs results from known motifs were used.

## Acknowledgements

We thank all the members of the NTase laboratory for helpful comments and discussion. We thank Frank Schwede (Biolog LSI) for his assistance in conceptualizing the nucleotide library. The work was funded by the German Research Foundation Emmy Noether Program 458004906 (C.C.O.M.), European Research Council ERC-StG-2023 101117085 (C.C.O.M.) and a FEBS Excellence Award (C.C.O.M.). I.B. is supported by a DFG Walter Benjamin Fellowship.

## Author Contributions

The project was conceived and experiments designed by I.B. and C.C.O.M. I.B. performed RNA-seq experiments and data analysis as well as live-cell imaging experiments. M.R.S. performed RT-qPCR analysis. RNA-seq library preparation and sequencing were performed by R.Ö. and R.R. The manuscript was written by I.B., and C.C.O.M., and all authors contributed to editing the manuscript and support the conclusions.

## Data availability

Raw sequencing data have been uploaded to European Nucleotide Archive (ENA) database.

---

**Figure S1.**
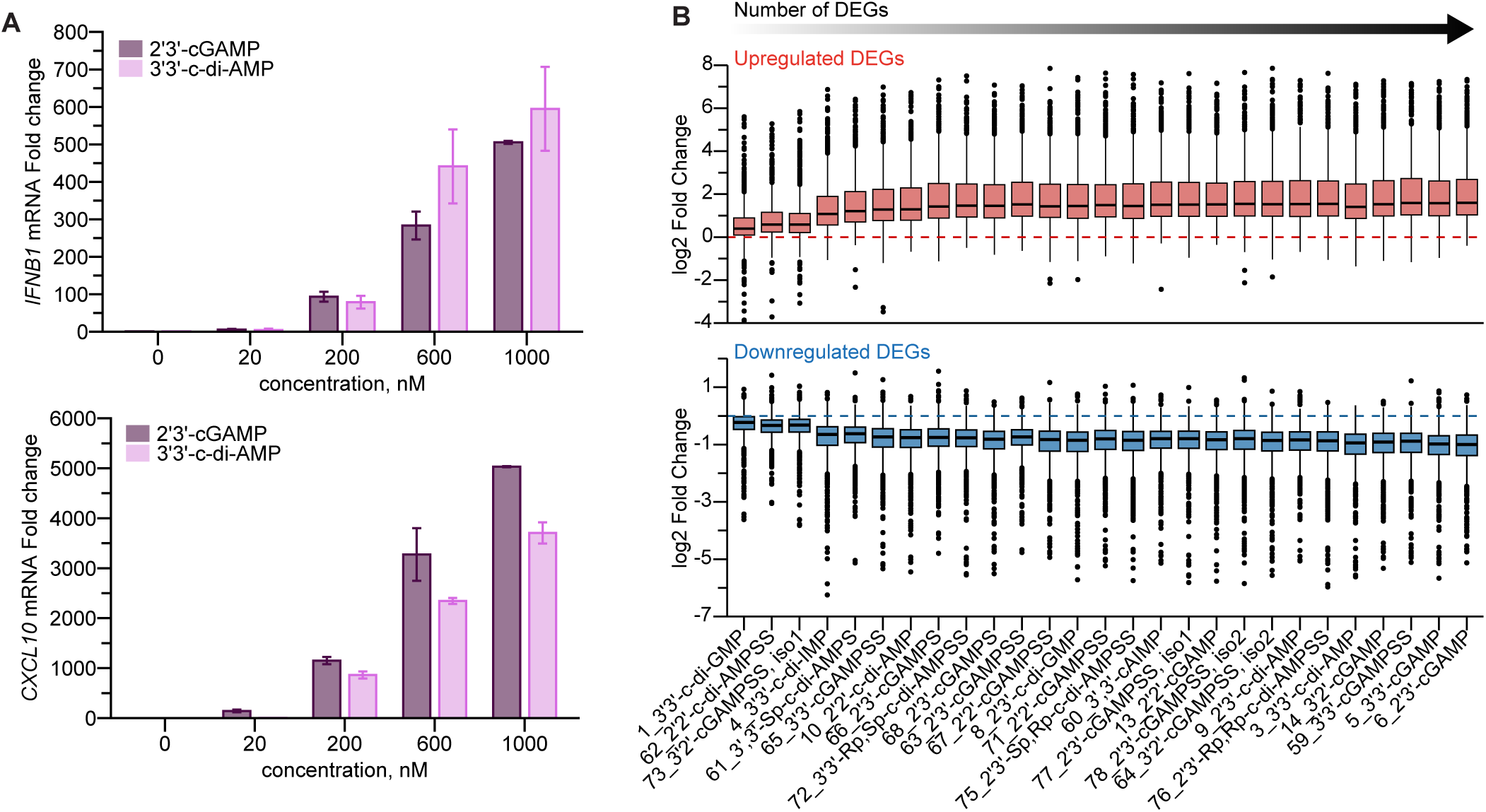
Analysis of nucleotide signaling in THP-1 cells, related to Fig 1. (A) RT-qPCR analysis of *IFNB1* and *CXCL10* expression after electroporation of THP-1 monocytes with indicated amount of 2′3′-cGAMP and 3′3′-c-di-AMP for 6 h. Bars represent means from two replicates and error bars represent standard deviation. (B) Boxplots showing log2 fold change for each CDN that induced signaling for upregulated and downregulated DEGs when pooling all DEGs from 27 signaling CDNs together. CDNs are sorted by the number of DEGs that they induced. Boxes encompass the twenty-fifth to seventy-fifth percentile changes. Whiskers extend to the tenth and ninetieth percentiles. Outliers are depicted with black dots. The central horizontal bar indicates the median.

**Figure S2.**
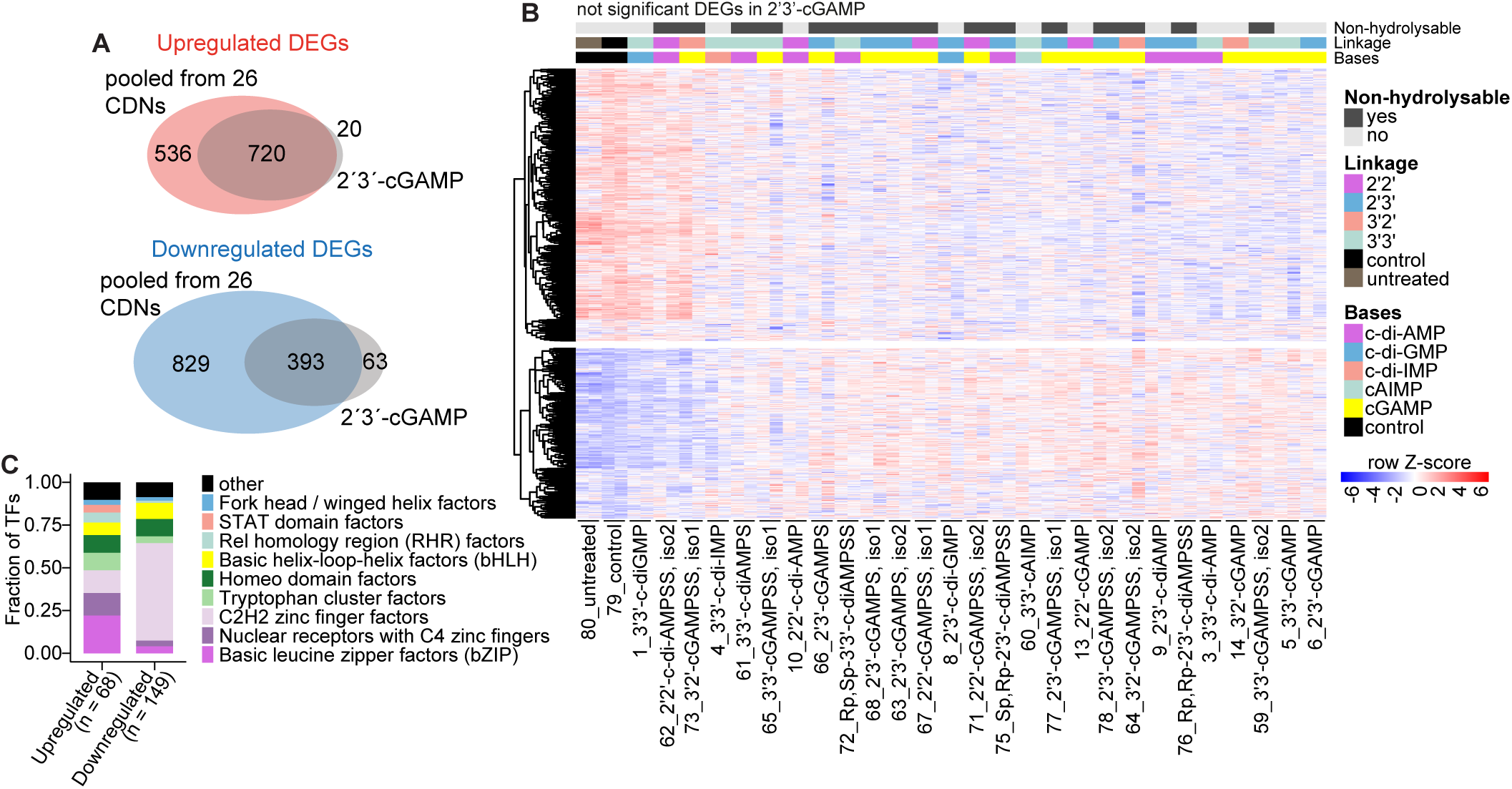
Analysis of gene expression changes that are induced by CDNs, related to Fig 2. (A) Overlap of upregulated and downregulated DEGs between 2′3′-cGAMP and all the DEGs pooled from the 26 other CDNs, which induced signaling. (B) Heatmap showing clustering of all DEGs from (A), which were not DEGs and were not significantly regulated by 2′3′-cGAMP (536 upregulated and 829 downregulated genes). Treatments are sorted by the number of DEGs, which they induced. Normalized transcript counts from variance stabilizing transformation (vst) were used, counts are row scaled. (C) Bar plot showing transcription factor families in the upregulated and downregulated genes from Fig 2C belonging to regulation of transcription based on the analysis with TFcheckpoint database.

**Figure S3.**
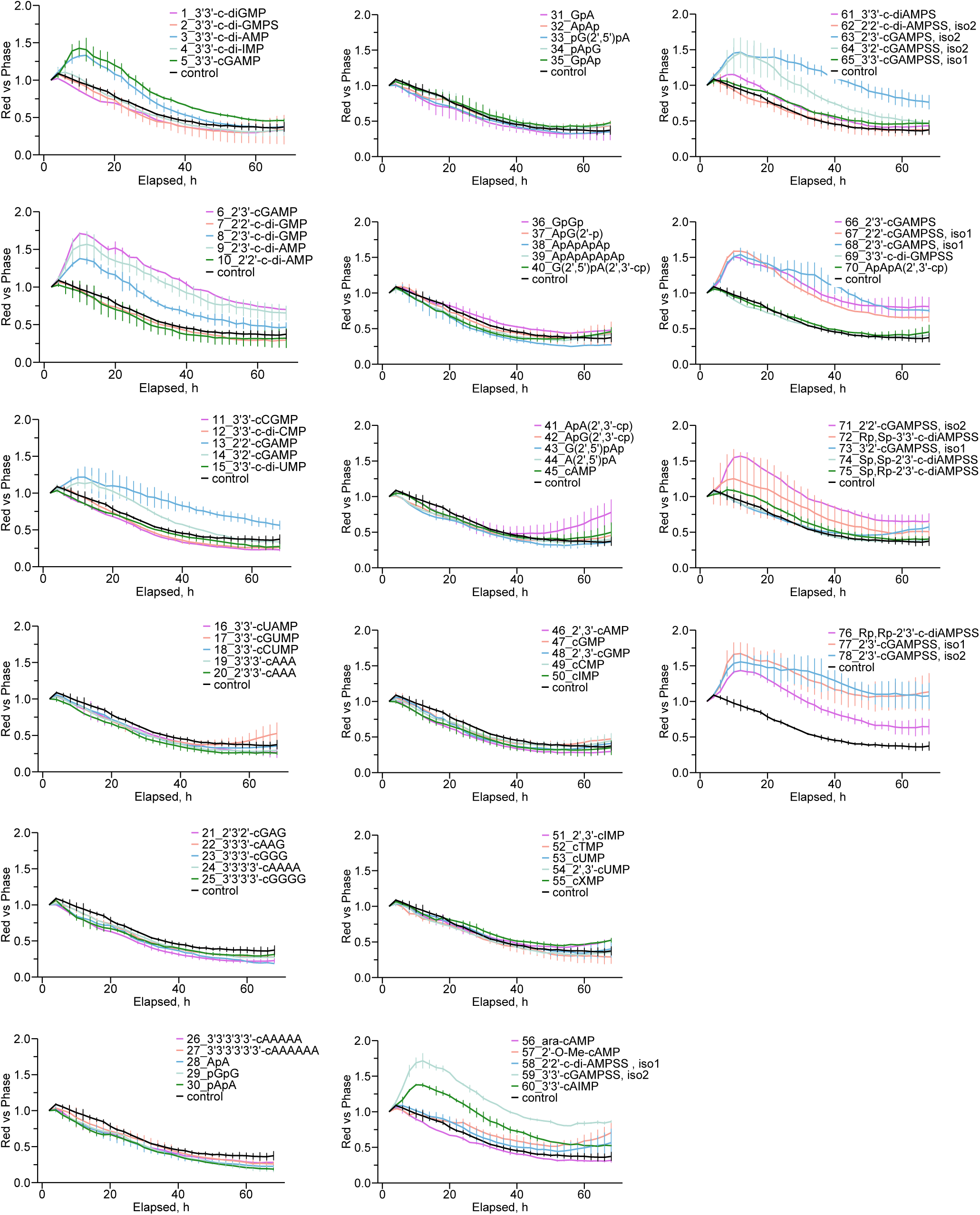
Analysis of nucleotide signaling in pancreatic cancer cells, related to Fig 4. Line plots showing induction of cell death for each nucleotide after electroporation of nucleotide library in human pancreatic cancer DANG cells. Cell death was quantified as red signal from YOYO-3 fluorescence vs phase. Lines represent means from two replicates, error bars indicate standard deviation. For all conditions red vs phase signal reduces over time due to cell attachment and recovery after electroporation.

**Table S1.**
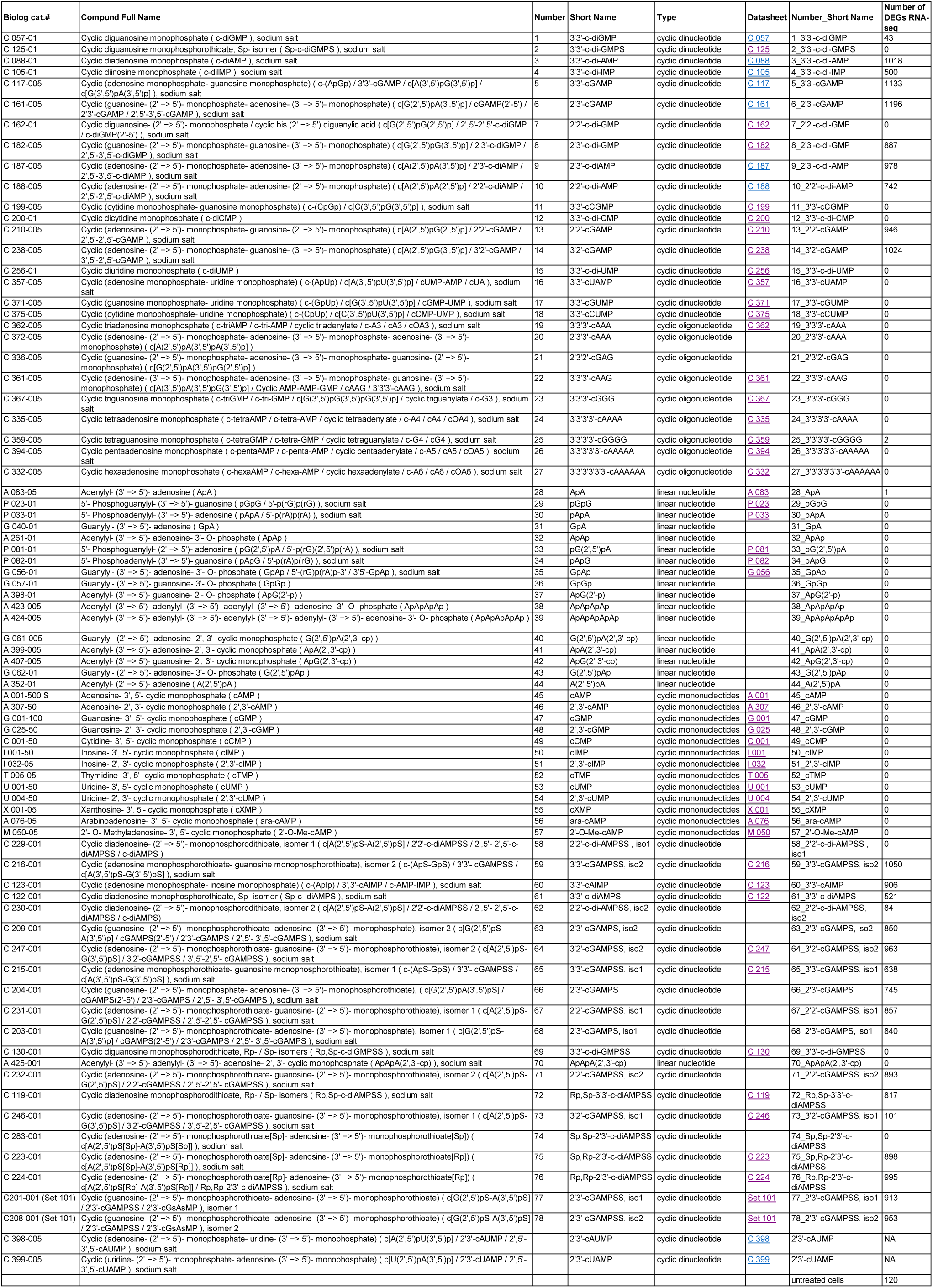
List of nucleotides in the nucleotide library and the number of DEGs identified from RNA-seq.

